# Elevated concentrations of polymyxin B elicit a biofilm-specific resistance mechanism in *Vibrio cholerae*

**DOI:** 10.1101/2023.06.26.546579

**Authors:** Julien Pauzé-Foixet, Marylise Duperthuy

## Abstract

*Vibrio cholerae* can form biofilms both in the aquatic environment and in the human intestine, facilitating the release of hyper-infectious aggregates. Due to the increasing antibiotic resistance that impedes treatment of infections, alternatives need to be found. One of these alternatives is antimicrobial peptides, including polymyxin B (PmB), which is already used to treat infections caused by antibiotic-resistant bacteria. In this study, we first investigated the resistance of *V. cholerae* O1 El Tor strain A1552 to various antimicrobials under aerobic and anaerobic conditions. An increased resistance to PmB is observed in anaerobiosis, with a 3-fold increase in the dose required for 50% growth inhibition. We then studied the impact of the PmB on the formation and the degradation of *V. cholerae* biofilms to PmB. Our results show that PmB affects more efficiently biofilm formation under anaerobic conditions. On the other hand, preformed biofilms are susceptible to degradation by PmB at concentrations close to the minimum inhibitory concentration (MIC), resulting in approximately 50% reduction of the biomass. At higher concentrations, we observed less degradation and an opacification of the biofilm structures within 20 minutes post-treatment, suggesting a densification of the structure. This densification does not seem to result from the overexpression of matrix genes but rather from the release of DNA through cellular lysis, forming a protective shield that limits the penetration of the PmB into the biofilm.

**Importance:** *Vibrio cholerae* is an intestinal pathogen capable of forming biofilms and resisting antimicrobials both in the aquatic environment and during infection. Understanding and determining the resistance of *V. cholerae* to antimicrobials during the infection is crucial to improve patient care. During the infection and in the aquatic environment, *V. cholerae* form biofilms, structures that are known for their significance in antimicrobial resistance. In this study, we investigated the antimicrobial resistance of *V. cholerae* in both aerobic and anaerobic conditions, in their planktonic and biofilm forms. The major finding of this study is the identification of a resistance mechanism specific to elevated concentrations of polymyxin B, a last-resort antimicrobial used in the treatment of infections caused by multidrug-resistant Gram-negative bacteria. This resistance mechanism likely involves the lysis of bacterial cells on the surface of the biofilm, resulting in the release of DNA that provides a protective shield against PmB for bacteria within the biofilm matrix.

## Introduction

*Vibrio cholerae* is a pathogenic Gram-negative aquatic bacterium found in marine environments, including estuaries and coastal waters. In its natural habitat, these bacteria can be found in the form of mobile planktonic cells and biofilms, sometimes in association with the chitin exoskeleton of zooplankton and crustaceans (1, 2). *V. cholerae* is also a facultative anaerobic bacterium with the ability to carry out both fermentation and anaerobic respiration (3). Anaerobic respiration requires the use of alternative electron acceptors such as Trimethylamine N-oxide (TMAO) to re-establish the electron transfer chain required for the synthesis of adenosine triphosphate (ATP) (3). *V. cholerae* is a pathogenic agent capable of infecting the human small intestine, where it can cause life-threatening episodes of diarrhea, clinically known as cholera. This illness is typically contracted by ingesting bacteria-contaminated food or water. Infected individuals excrete high levels of both planktonic bacteria and biofilm-like aggregates during episodes of diarrhea, which further contribute to the fecal-oral infectious cycle (2). The formation of biofilm-like aggregates during the infection of the gut improves host infectivity by increasing the resistance of bacteria to stresses, during both the infection and the excretion of infectious particles into the surrounding environment (4). Populations with no access to clean drinking water are most at risk of contracting cholera in endemic developing countries and during epidemics triggered by humanitarian disasters. (5).

Biofilms are tridimensional structures made up of bacterial aggregates embedded in a matrix primarily composed of polymers, proteins and nucleic acids (6). The biofilm lifestyle brings several benefits to bacteria, including enhanced environmental persistence, improved antimicrobial resistance, an optimal setting for horizontal gene transfer and social cooperation through quorum sensing. (7). Due to their greater resistance and persistence on both organic and abiotic surfaces, biofilms pose an additional challenge to the control of bacterial infections, notably in terms of sanitation in healthcare facilities (8, 9). According to the NIH, over 60% of all bacterial infections and around 80% of chronic infections involve biofilms (10).

In *V. cholerae*, the biofilm’s properties include improved resistance to gastric acid and a greater competitive edge over nutrients in the small intestine (11). The vibrio exopolysaccharide (VPS) matrix is the product of the *vps* genes, which are split into two operons: *vps-I* and *vps-II*. (12). On the *V. cholerae* chromosome I, the *vps* operons are spatially separated by the *rbmA-F* genes, which mostly code for proteins involved in biofilm structure. RbmA protein promotes adhesion between VPS-producing bacteria and recruitment of free-living planktonic bacteria (13). RbmC protein plays a stabilizing role in the biofilm structure and interacts with Bap1 to build a protein matrix that surrounds cell aggregates (14). Bap1 also enables biofilm to adhere to its substrate (15). Besides proteins and VPS, *V. cholerae*’s biofilm matrix also contains extracellular DNA (eDNA). The level of eDNA is controlled by the Dna and Xds proteases that have a role in the development of a typical three-dimensional biofilm structure, detachment from a mature biofilm and utilization of eDNA as a nutrient source (16). *V. cholerae* can take the form of rough variants when the environmental conditions are unfavorable (17). These variants produce greater volumes of biofilm, which provides them with improved resistance to both osmotic and oxidative stresses, thus contributing to bacterial persistence and further dissemination in human populations. (18). Rough variants are more common in clinical strains than in the environmental isolates of *V. cholerae*, thus representing a potentially important evolutive and adaptive benefit (19).

The enhanced antimicrobial resistance of biofilms is largely due to the impaired ability of antimicrobial molecules to diffuse throughout the biofilm matrix, the increased expression of antibiotic resistance genes induced by the quorum sensing process, and the coexistence of different metabolic stages in the bacterial community (20). Antimicrobial peptides (AMPs) have a broad spectrum of activities, including the destabilization of the outer membrane of Gram-negative bacteria, and interacting with intracellular targets (21). In addition to their bactericidal activities, the subinhibitory concentrations of AMPs can have an impact on a variety of *V. cholerae* physiological functions, such as virulence and inhibition of biofilm production (22, 23). In contrast, in some other species, biofilm production can be enhanced by sub-inhibiting levels of Polymyxin B (PmB), a last-line antimicrobial peptide used to counter antibiotic-resistant bacteria infections (24). Among the AMPs, the polymyxins, including the polymxin B (PmB), are already used as last line antimicrobial to treat infection caused by multidrug resistant Gram-negative pathogens in human and animal health (25).

In this study, we first investigated the impact of anaerobiosis on antimicrobial resistance using TMAO as alternative electron acceptor. We then evaluated the impact of subinhibitory concentrations of PmB on biofilm formation under aerobic and anaerobic conditions. Finally, we determined the impact of PmB on mature biofilm and found a non-linear dose-dependent effect on the biofilm biomass, with a maximal reduction of biomass at concentrations close to the minimal inhibitory concentration (MIC). Above this concentration, the biofilm appears to be more resistant to PmB, which might be due to the massive release of DNA.

## Material and methods

### Bacterial Strains and growth conditions

All strains and plasmids used in this study are listed in Table I. *V. cholerae* O1 El Tor Inaba A1552 is a clinical strain isolated from a Peruvian traveller in the 1990s during an outbreak of cholera in South America (26). Strain A1552R is a rough variant phenotype that produces larger amounts of biofilm (27). Bacteria were grown in lysogeny broth (LB) medium (10 mg/ml tryptone (Termo Fisher™), 5 mg/ml yeast extract (Thermo Fisher™), 5 mg/ml NaCl). LB supplemented with 50mM TMAO is used to promote anaerobic respiration in anaerobic conditions (28). Experiments under anaerobic conditions were performed in a Bactron 600 anaerobic chamber (Sheldon Manufacturing) equipped with an incubator. All media intended for anaerobic growth were stored in the anaerobic chamber for 48 hours prior to experimentations, to ensure removal of any residual oxygen. All experiments were performed at a constant temperature of 37°C.

**Table I:**
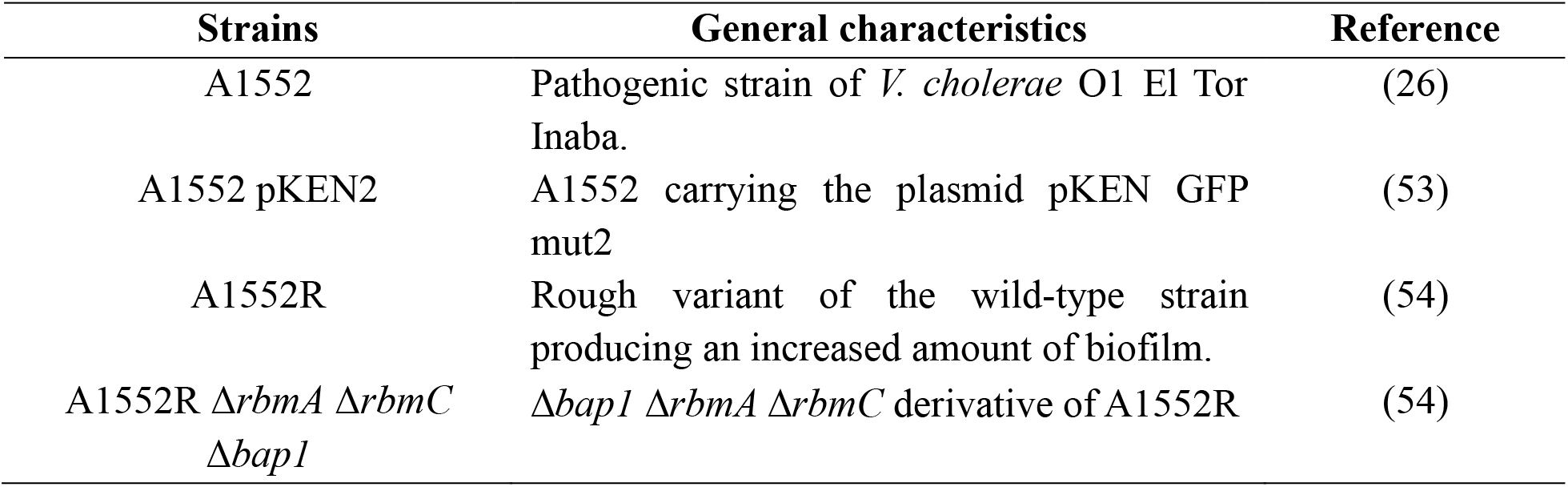
Strains used in this study

| Strains | General characteristics | Reference |
| --- | --- | --- |
| A1552 | Pathogenic strain of *V. cholerae* O1 El Tor Inaba. | (26) |
| A1552 pKEN2 | A1552 carrying the plasmid pKEN GFP mut2 | (53) |
| A1552R | Rough variant of the wild-type strain producing an increased amount of biofilm. | (54) |
| A1552R $\Delta rbmA$ $\Delta rbmC$ $\Delta bapI$ | $\Delta bapI$ $\Delta rbmA$ $\Delta rbmC$ derivative of A1552R | (54) |

### Determination of the minimum inhibitory concentration (MIC) and the half maximal inhibitory concentration (IC_50_)

The minimum inhibitory concentrations were determined as previously described (29) in LB or LB-TMAO. Plates were incubated at 37°C with shaking in aerobic condition, or without agitation in anaerobic condition, for 18h. The optical density at 600nm (OD_600nm_) was measured using SpectraMax® ID3 plate reader (Molecular Devices). The minimum inhibitory concentration (MIC) is defined as the smallest concentration that inhibits bacterial growth to less than 10% of their maximal growth. The half maximal inhibitory concentration (IC_50_) is the lowest concentration that inhibits the bacterial growth to less than 50% of their maximal growth. All assays were performed in biological and technical triplicates.

### Impact of antimicrobials on biofilm formation and degradation

Bacteria were grown in LB at 37°C under shaking at 250 RPM overnight, and then diluted [1:100] in fresh LB. 100 µl of the diluted culture were distributed in a 96 well-plate and incubated at 37°C without shaking for 18 hours. The wells were drained and rinsed with distilled water. Biofilms were then stained as previously described (30). Briefly, 100 mL of a solution of 0.1% crystal violet was added to the well. After 10 minutes, the wells were rinsed with distilled water and allowed to dry for 24 hours. The biofilms were dissolved in 100 µl of 30% acetic acid and the optical density at 595 nm (OD_595nm_) was measured using a SpectraMax® ID3 plate reader (Molecular Devices) to quantify the biofilm biomass. A standardized value was removed for each PmB concentration to blank the samples. These values were determined by incubating 100 µl of LB with various concentrations of PmB that have been subjected to the same staining steps as the biofilms. In biofilm formation assays, PmB was added to 96-well plates at the beginning of the bacterial growth. In biofilm degradation assays, PmB was added to a 24-hours preformed biofilm, which was further incubated for 24 hours before staining. All assays were repeated in biological and technical triplicates.

### Microfluidic microscopy

1. *V. cholerae* biofilm growth was monitored using the microfluidic system Bioflux200 (Fluxion) using the A1552 pKEN2. The microfluidic system was set up according to the manufacturer’s recommendations. The capillaries were inoculated with 20 µl of bacterial culture at an OD_600nm_ of 0.5, and then incubated for 30 minutes to allow the bacteria to adhere. After cleansing the planktonic cells, the system was launched with a flux of 0.20 dynes. The stream was left running continuously under a heated plate at 37°C for the duration of data acquisition. The effect of PmB on biofilm formation was determined by the addition of PmB to the culture media after the initial adhesion step. The effect of PmB on preformed biofilm was determined 12h after the initial adhesion step by washing the biofilm and replace the culture media by PmB-supplemented LB. For the survival assays, the wells were rinsed after flow-through with PmB during 2 hours and replaced with fresh LB. After a 5-minute wash cycle at 1 dyne and 1h at regular flow, a final rinse was performed to eliminate any residual PmB. Pictures were captured with a Revolve4 microscope (Echo), in inverted mode, at 100x magnification (Olympus Objective 10X Fluorite ELWD NA 3.0-4.2MM) in bright field. Fluorescence images were taken using a DAPI filter for NucBlue™ and a Texas Red filter for propidium iodide (PI).

### Biofilm formation on solid media

Sterile cellulose filters (0.22 µm) were deposited on LB agar plates and 10 µl of *V. cholerae* A1552 or A1552R cultures were spotted at their center. After overnight incubation at 37°C, the filters were transferred to LB agar plates supplemented with increasing concentrations of PmB. After 72 hours of growth, colony phenotypes were observed.

### Gene expression

*V. cholerae* A1552R biofilms were grown in 24-well plates for 24 hours. The medium was then replaced by fresh medium with PmB when required and incubated for 15 minutes or 6 hours. The medium and surface pellicle were then removed. Three wells with the same culture conditions were treated with 333µl of TRIzol™ and pooled to obtain a total volume of 999µl per experimental condition. After RNA extraction according to the manufacturer’s instructions, the RNA was treated using the TURBO DNA-free Kit (Invitrogen). cDNAs were synthesized using the High-Capacity cDNA Reverse Transcription Kit (AppliedBiosystems). Quantitative PCR (qPCR) were carried out as previously described (31) using the primers listed in table II. The expression of each gene was calculated in PmB treated bacteria in comparison to non-treated cells using QuantStudio™ Desing and Analysis Software (Thermo FisherTM, Waltham, USA) v1.5.1 and normalized using *recA*. At least 4 biological replicates and 3 technical replicates are used to generate results.

**Table II:**
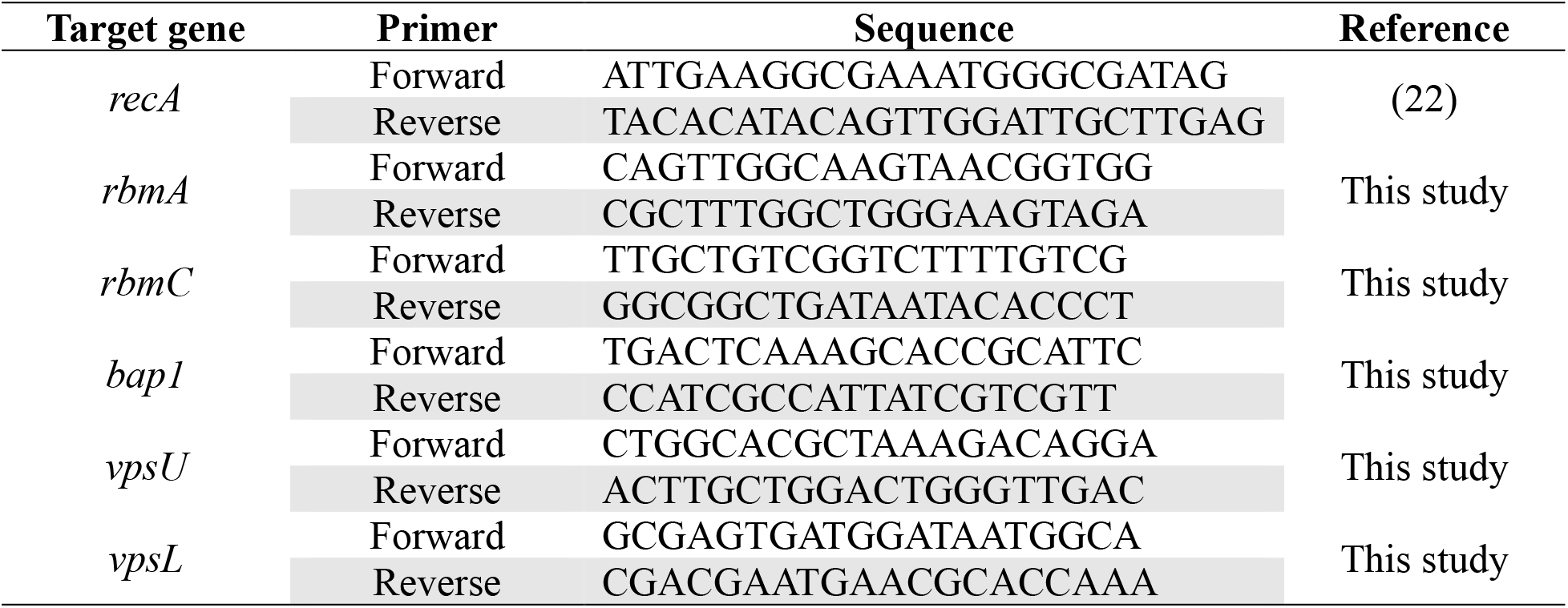
Primers used in this study

## Results

### Resistance of *V. cholerae* to PmB increases in anaerobiosis

To determine whether the antimicrobial resistance of *V. cholerae* O1 El Tor is affected by the oxygen availability in its environment, we conducted antibiotic resistance tests in the presence and absence of oxygen. Aerobic tests were performed in LB medium with agitation for 18 hours, while anaerobic tests were carried out in LB medium supplemented with 50mM TMAO as the final electron acceptor, without agitation in an anaerobic chamber. Different antimicrobials were tested, and the minimum inhibitory concentration (MIC) and the half maximal inhibitory concentration (IC_50_) for each antimicrobial were determined for both A1552 and A1552R strains.

There are no differences between the MICs and IC_50_ of A1552 and A1552R variant (Table III). The maximum growth of both A1552 and A1552R is reduced under anaerobic conditions in comparison to aerobic growth (Figure 1). There is an increased resistance to the antibiotics belonging to the aminoglycoside family, namely streptomycin (Strep) and kanamycin (Kan) under anaerobic conditions, with MICs and IC_50_ values that are two times higher than in aerobic conditions (Table III). No differences were observed for chloramphenicol, carbenicillin, azithromycin, erythromycin, tetracycline, doxycycline, and ciprofloxacin, both in terms of MIC and IC_50_ (Table III). As for the PmB, given the high resistance of *V. cholerae* O1 El Tor to it, we tested the effect of concentrations from 25µg/ml to 250 µg/ml on bacterial growth (Figure 1). The MICs are the same in aerobic and anaerobic conditions (Table III). Interestingly, there is a significant increase of the IC_50_ under anaerobic conditions, which is three times higher than under aerobic conditions (Table III) (Figure 1). It is also noteworthy that the growth of the A1552R variant appears to be stimulated by low concentrations of PmB when compared to the control without PmB (Figure 1).

**Figure 1:**
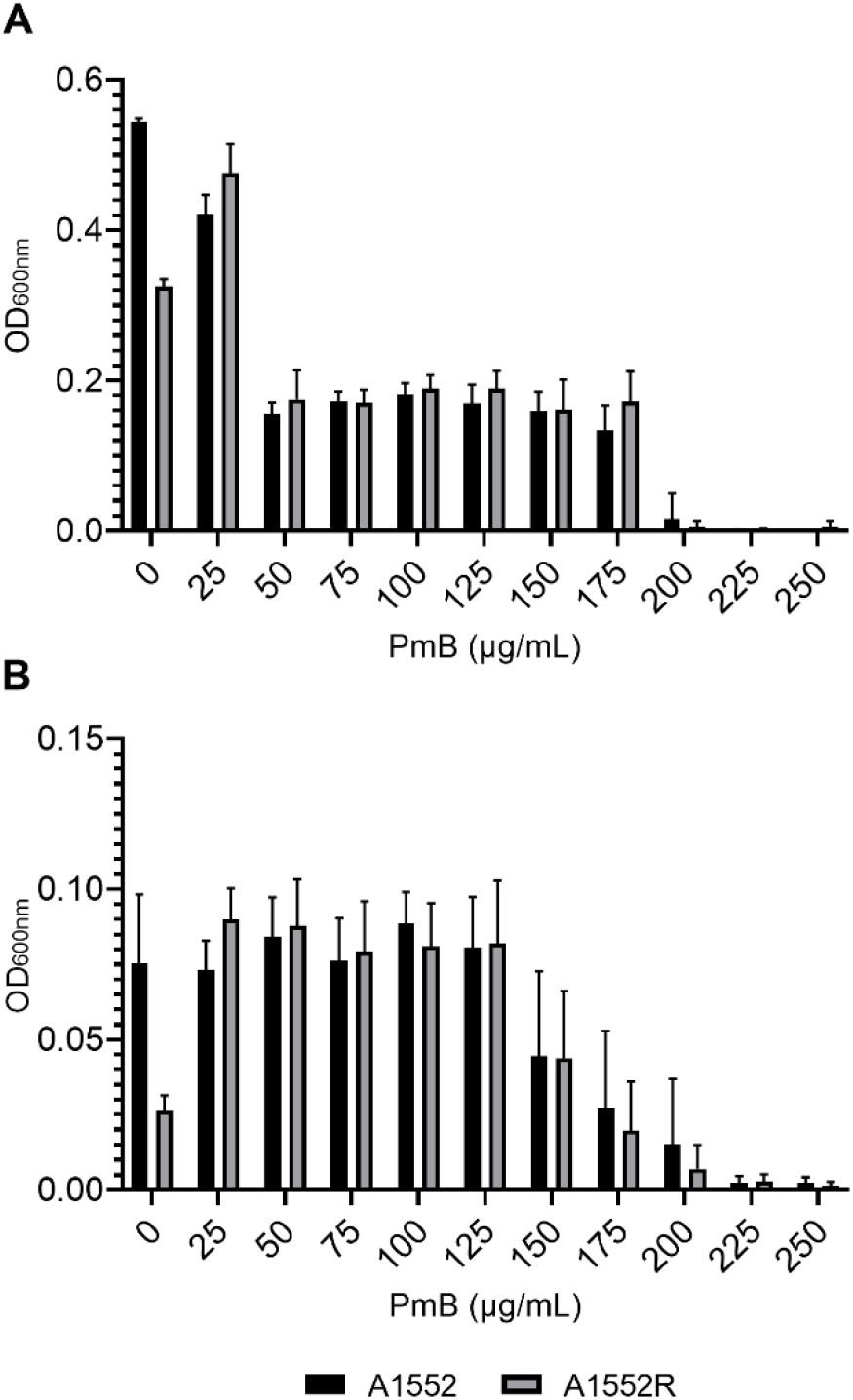
Growth of *V. cholerae* A1552 and A1552R under different concentrations of PmB in aerobic (A) and anaerobic (B) conditions. The bacteria were grown with or without PmB in 96- well plates at 37°C and the OD_600nm_ was measured after 18h. Data represent mean ± standard deviation (SD) of three independent experiments conducted in triplicates.

**Table III:**
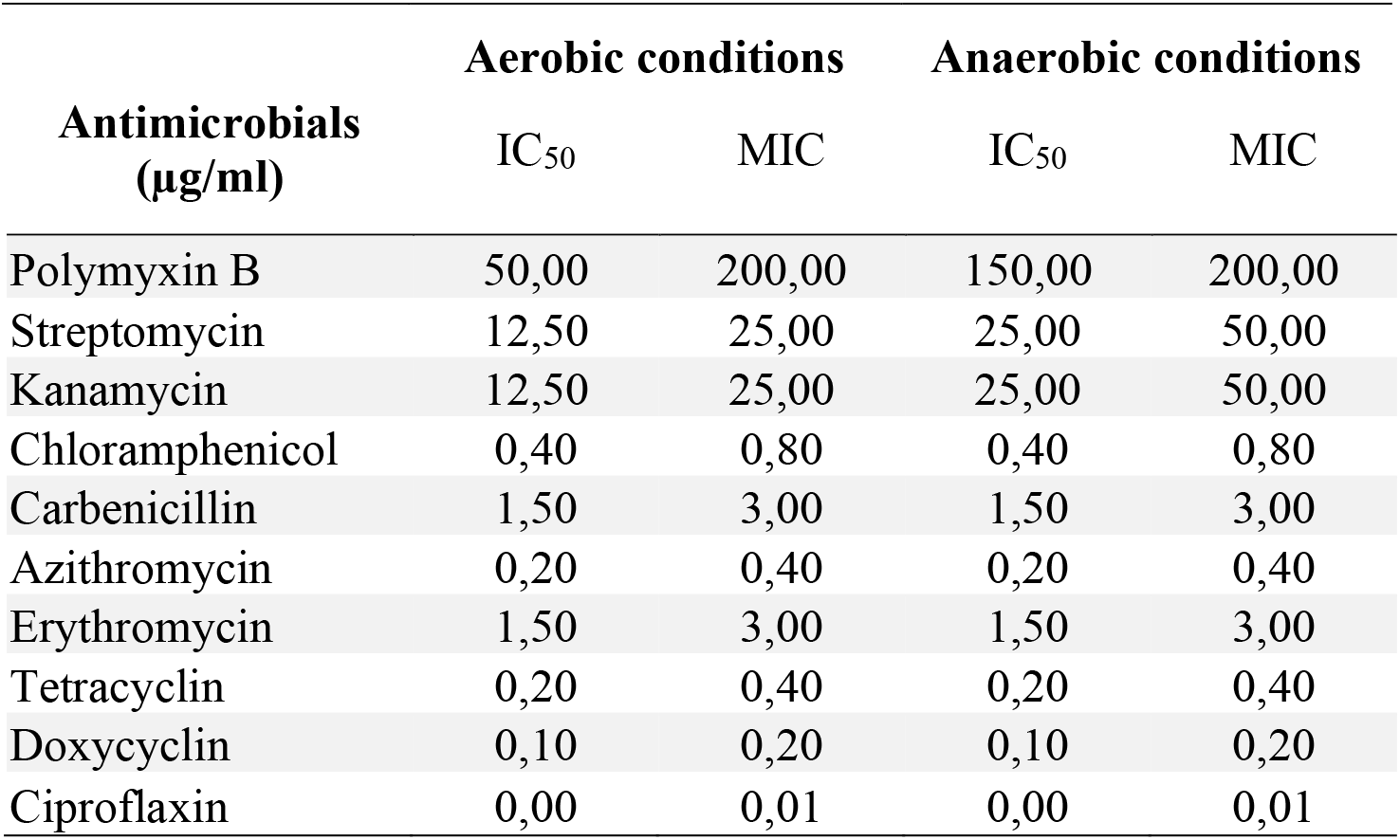
Minimal inhibitory concentration (MIC) and half maximal inhibitory concentration (IC_50_) in A1552 and A1552R determined under aerobic and anaerobic conditions.

### The reduction of biofilm formation in the presence of PmB is further enhanced under anaerobic conditions

Given the importance of biofilms in antimicrobial resistance and because of the increased resistance toward PmB in anaerobiosis, we investigated the biofilm production in the presence of low concentrations of PmB. To evaluate biofilm production, we used the A1552R and A1552 strains, as well as the triple mutant A1552RΔ*rbmA*Δ*rbmC*Δ*bap1*, lacking the essential matrix proteins as a negative control. Biofilm production assays were conducted in LB medium at 37°C without agitation for 18 hours, with or without PmB, and biofilm biomass was quantified using crystal violet. As expected, in the absence of PmB, the A1552R strain produces a significantly higher biofilm biomass compared to the A1552 strain, as demonstrated by a higher OD_595nm_, while the *rbmA*-*rbmC*-*bap1* triple mutant has a severely impaired ability to form a biofilm (Figure 2AB). Furthermore, the biofilm formation was almost entirely abolished at 25 µg/ml and 3 µg/ml under aerobic and anaerobic conditions, respectively (Figure 2AB). At lower concentrations, up to 6 µg/ml and 1.5 µg/ml under aerobic and anaerobic conditions, respectively, a less severe decrease in biofilm formation is observed (Figure 2AB). These results collectively indicate that the presence of PmB in the medium impairs biofilm formation at concentrations well below the MIC, and the absence of oxygen exacerbates this phenomenon despite the higher IC_50_.

### The efficiency of PmB to degrade mature biofilms of *V. cholerae* decreases at high concentrations

We then investigated the resistance of mature biofilms to degradation by PmB (Figure 2CD). Mature biofilms were treated with different concentrations of PmB for 24h and the biomass was quantified using crystal violet staining (Figure 2CD). Due to the highly resistant nature of biofilms, significantly higher concentrations of PmB were used, that were up to 1600 µg/ml. Control wells with PmB alone were stained with crystal violet to remove background noise caused by PmB adhering to the wells. In A1552, a decrease in biofilm biomass was observed, reaching a minimal at 25-50 µg/ml in aerobic condition compared to the non-treated biofilm (Figure 2C). Subsequently, an increase in biofilm biomass was gradually observed with increasing PmB concentrations, up to the highest concentration tested of 1600 µg/ml (Figure 2C). In anaerobic condition, we observe a decrease in the biofilm biomass (figure 2D). In the A1552R strain, an increase in biofilm biomass was observed at sub-inhibitory concentrations of PmB, specifically between 3 and 25 µg/ml under aerobic conditions (Figure 2C). This stimulation was not observed under anaerobic conditions (Figure 2D). The most significant decrease of biomass quantities was observed at 200 µg/ml, which represents the MIC of planktonic cells, and 50 µg/ml under aerobic and anaerobic conditions, respectively (Figure 2CD). Afterward, the biofilm biomass gradually increased with the PmB concentration in both conditions (Figure 2CD). These results suggest an increased resistance of the biofilm to degradation at high concentrations of PmB. Since we observed no major difference in absence and in the presence of oxygen, we pursued our investigations in aerobic conditions.

**Figure 2:**
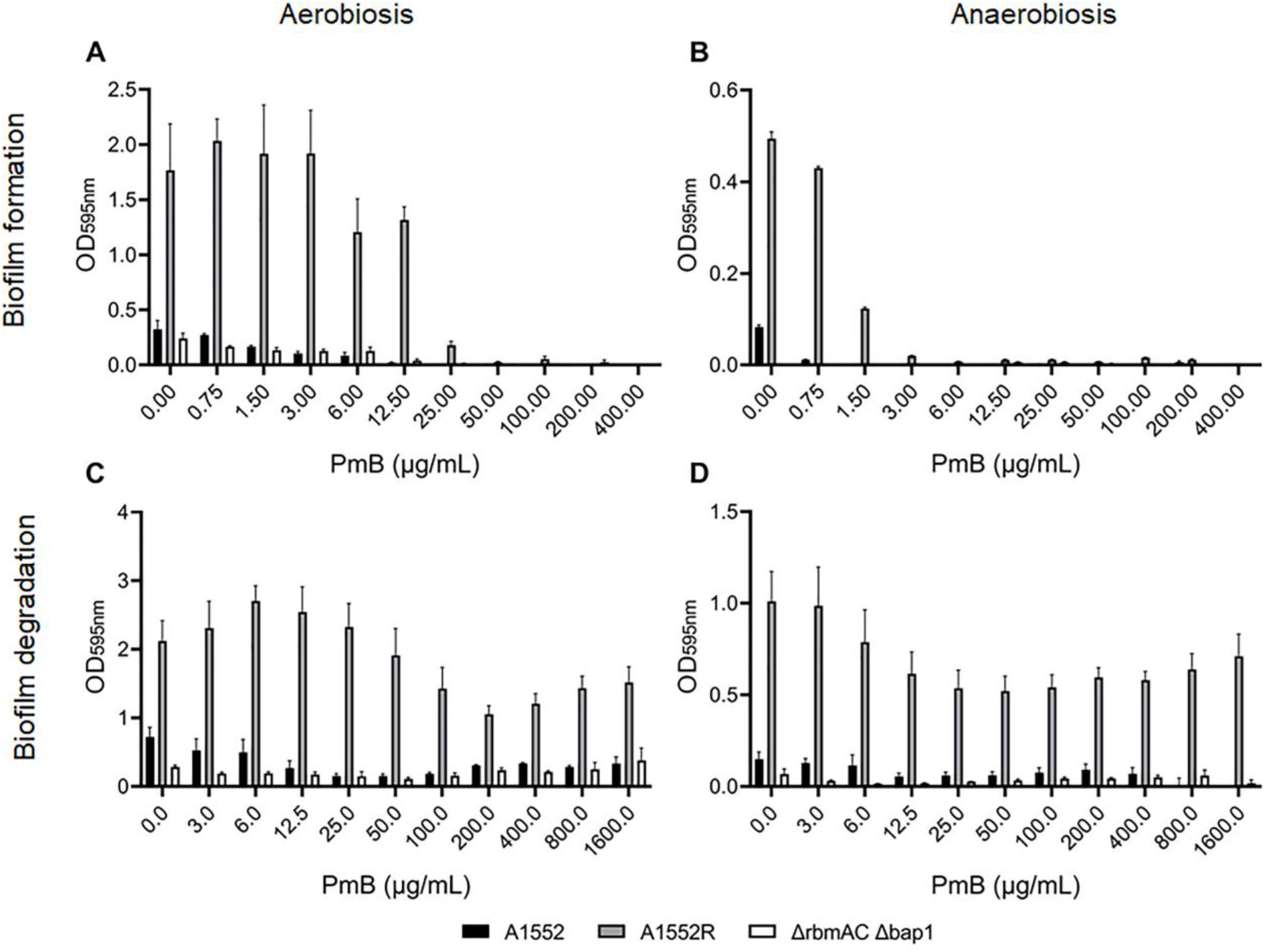
Impact of polymyxin B (PmB) on biofilm production (A & B) and biofilm degradation (C & D) in *V. cholerae* A1552, A1552R and A1552RΔ*rbmA*Δ*rbmC*Δ*bap1* in aerobiosis (A & C) and anaerobiosis (B & D) conditions. For the biofilm production, PmB was added to the culture media prior to biofilm formation for 18 h, whereas for the biofilm degradation, it was added on pre-formed biofilms and quantified after 24h of treatment. The biofilm biomass was quantified 24h after the PmB treatment using a crystal violet staining and a measurement of the OD_595nm_. Data represent mean ± 95% Confidence Intreval (CI) of three independent experiments conducted in triplicates.

Microscopic observations of biofilms were performed using the A1552R strain under bright field with and without crystal violet staining (Figure 3). We tested three concentrations of PmB: 6 µg/ml (PmB6), 200 µg/ml (PmB200), and 1600 µg/ml (PmB1600), representing the highest stimulation, reduction, and resistance of the biofilm, respectively. A control without PmB was included. Biofilm structures were observed at all concentrations, confirming that even at high concentrations, PmB is incapable of eradicating completely the biofilm (Figure 3). When treated with 6 µg/ml, the biofilm showed no major changes compared to the control without PmB (Figure 3). As expected, we observed a decrease in biomass at 200 µg/ml (Figure 3). In the biofilm treated with PmB1600, an increase in well coverage and staining intensity was observed, confirming an increase in biofilm biomass.

**Figure 3:**
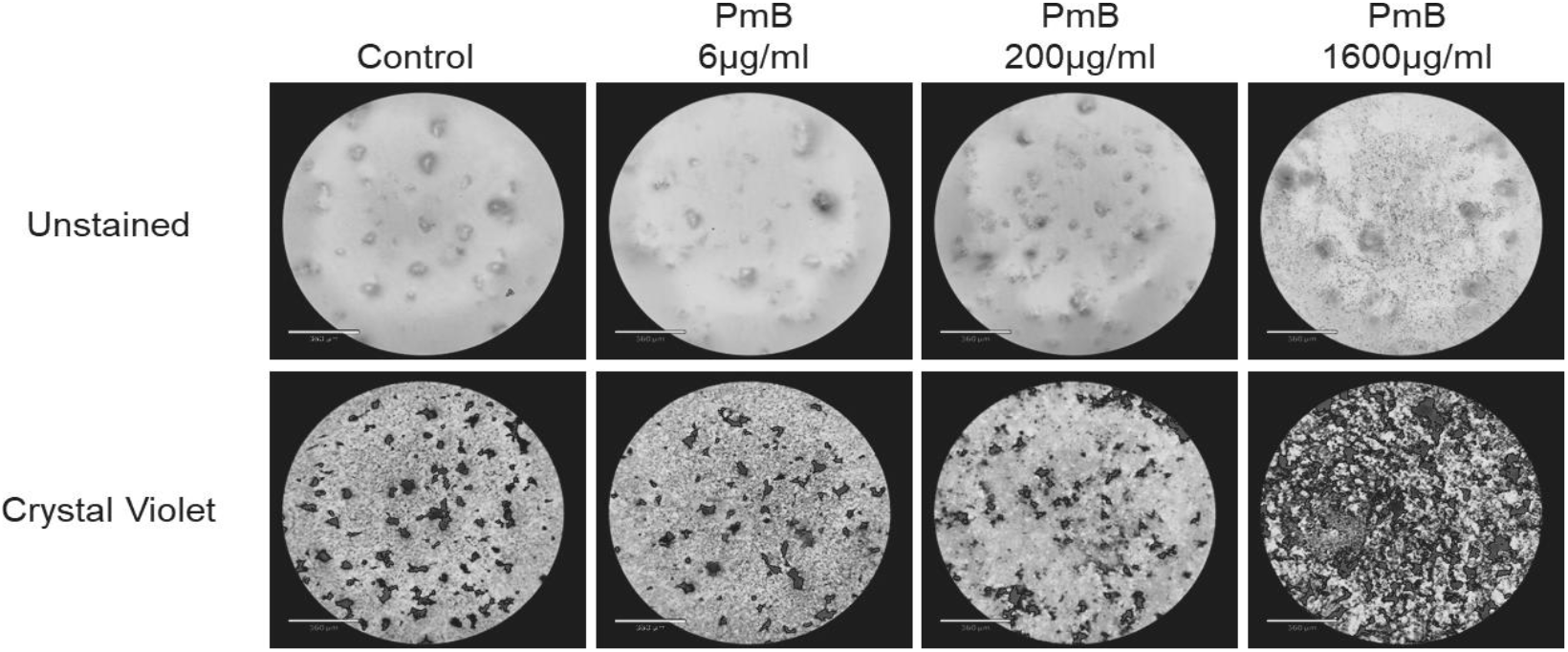
Preformed A1552R biofilms treated with polymyxin B (PmB) for 24h. Microscopic observations of unstained (upper panel) and crystal violet-stained (lower panel) biofilms were performed after a treatment with PmB for 24 hours in aerobiosis.

### High concentration of PmB induces the emergence of three-dimensional spike-like structures on solid media

To assess the impact of PmB on biofilm formation on solid medium, we exposed pre-formed colonies of *V. cholerae* to PmB (Figure 4). To do this, we placed 0.22 µm filters onto LB agar plates and spotted 10 µl of *V. cholerae* A1552 and A1552R cultures on top of them. Filters allow *V. cholerae* to use the nutrients from the media, but retain the bacteria on their top. After overnight incubation, the filters were transferred onto new LB agar plates supplemented or not with different concentrations of PmB (6, 200 and 1600 µg/ml). The plates were incubated for 3 days at 37°C. Colony morphology on PmB6 and on PmB200 appeared to exhibit a slightly increased roughness in both strains (Figure 4). Both strains grew at these concentrations, as determined by a wider colony diameter, albeit the growth on PmB200 was reduced compared to the non-treated control (Figure 4). On PmB1600, both strains displayed a distinct phenotype, with colonies not expanding and with the emergence of three-dimensional spike-like structures, suggesting a vertical expansion instead of a horizontal one.

**Figure 4:**
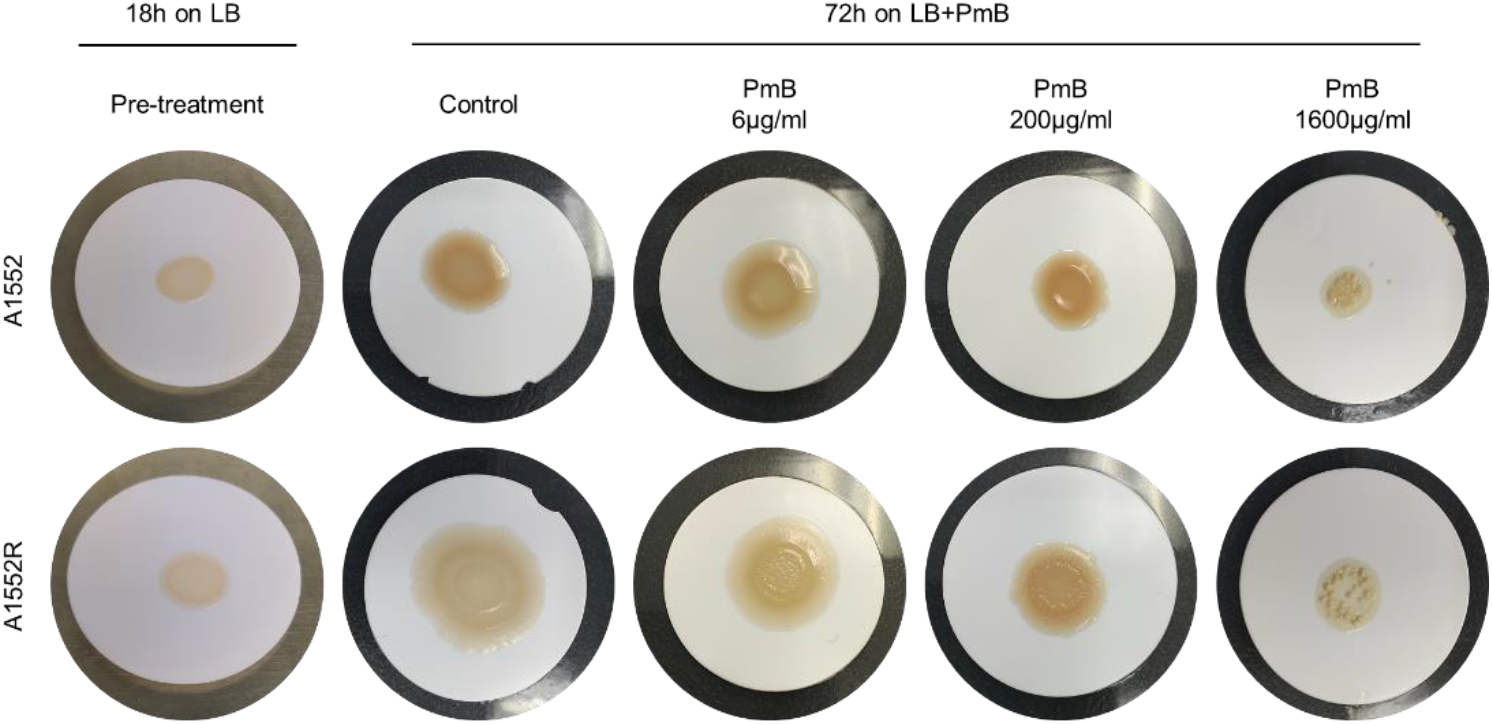
Impact of various concentrations of polymyxin B (PmB) on *V. cholerae* colony morphology. A droplet of *V. cholerae* A1552 (upper panel) and A1552R (lower panel) cultures was spotted on 0.22 μm cellulose filters placed on LB agar medium and incubated for 24 hours. The membrane was then transferred to an LB agar medium containing PmB and incubated for an additional 24 hours. The colony appearance was subsequently observed. Data are representative of at least 3 independent experiments.

### PmB induces the formation of poorly adherent biofilms

To better characterize *V. cholerae* biofilms formation in the presence of PmB, we transitioned to a microfluidic system and observed biofilm development. The system allows visualization of biofilm formation within a capillary under constant flow of medium. We used the A1552 pKEN2 strain for these observations, as the production of a GFP protein helps to visualize bacteria in the system. After an initial 30-minute adhesion of bacteria, the biofilm was allowed to form within the capillary under flow of LB medium with or without PmB, providing a continuous supply of nutrients at 37°C. Images were captured at regular intervals to characterize biofilm development (Figure 5).

**Figure 5:**
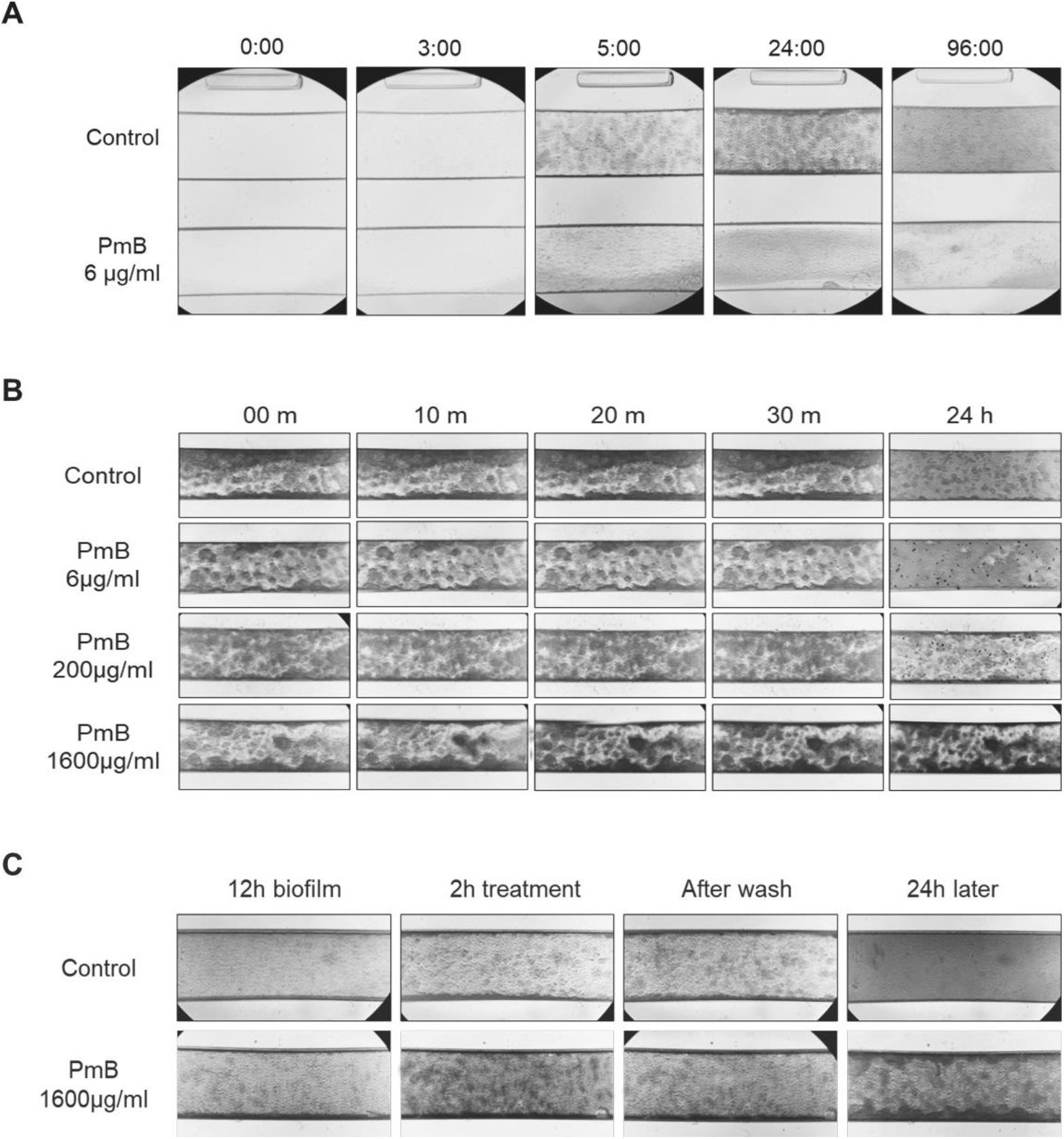
Impact of polymyxin B (PmB) on A1552 biofilm formation (A), biofilm degradation (B) and survival (C) in a microfluidic system. A) To assess the impact of PmB on biofilm formation, PmB at 6, 200 and 1600 μg/ml was added to the culture media right after the initial adhesion step, and biofilm formation was monitored over a period of 96 hours. B) The degradation of the biofilm by PmB was determined by adding PmB at 6, 200 and 1600 μg/ml to a 12-hour preformed biofilm. The impact on the biofilm was assessed at 10, 20, 30 minutes, and 24 hours. C) A 12-hour preformed biofilm was treated with 1600 μg/ml of PmB for 2 hours, washed, and then incubated without PmB for 24 hours to assess bacterial survival. Data are representative of at least 3 independent experiments.

In the control condition without PmB, we observed biofilm formation as early as 3 hours post-adhesion, which further developed into dense structures after 5 hours (Figure 5A). By the fourth day, no flow was visible, indicating that the capillary was clogged by the biofilm. In the presence of low concentrations of PmB (PmB6), we observed a significantly slower development of the biofilm (Figure 5A). In addition to producing less biofilm, it appeared less firmly attached to the capillary wall, as observed at the 24-hour time point, and less structured with the absence of dense structures (Figure 5A). Even after 4 days of flow, the capillary containing PmB still exhibited movement. These results suggest that low concentrations of PmB in a dynamic system under medium flow induce a decrease in biofilm adhesion to the substrate and a loss of biofilm structuring.

### A treatment of pre-formed biofilm with PmB quickly induces the densification of the mature structures

We then tested the impact of PmB on pre-formed biofilms in the microfluidic system (Figure 5B). We initiated flow in absence of PmB until a biofilm was established (12 hours), while still allowing flow circulation. Subsequently, we changed the medium and added different concentrations of PmB (6, 200, and 1600 mg/ml). The PmB-containing medium was then left to flow at 37°C for 24 hours.

There was very little difference observed within the first 30 minutes for the control, PmB6 and PmB200 concentrations (Figure 5B). However, PmB1600 showed a noticeable difference as early as the first 20 minutes of contact with the PmB, and at 20 minutes, the biofilm appeared significantly darker than in the non-treated control (Figure 5B). Between 30 minutes and 24 hours, no visible difference was observed for PmB1600, suggesting that the change occurs rapidly, predominantly within the first 20 minutes of treatment (Figure 5B). Additionally, in PmB1600 condition, the biofilm did not develop in the uncovered regions of the capillary as no new structures were observed there (Figure 5B). Conversely, the non-treated control eventually covered the entire capillary within 24 hours, while PmB6, although more developed than initially, did not cover the entire capillary (Figure 5B). In agreement with the biomass results determined by crystal violet staining (Figure 2), PmB200 was the only concentration that showed less biofilm after 24 hours of exposure to PmB, indicating that a portion of the biofilm was disaggregated by PmB. Altogether, our results suggest that a treatment with PmB on pre-formed biofilms has contrasting effects depending on the concentration. At concentrations of 200 µg/ml, which represents the MIC of the planktonic form, the biofilm is reduced, while at higher concentrations (1600 µg/ml), the biofilm appears denser through a phenomenon that occurs within the first 20 minutes of treatment and persists over time (Figure 3,5).

Given that the concentration of 1600 µg/ml is 8 times higher than the MIC and the absence of new structures after 24 hours of treatment, we wondered if the biofilm embedded bacteria were still alive (Figure 5C). To assess this, a 12-hour mature biofilm under flow was treated with 1600 µg/ml of PmB. Knowing that PmB acts quickly on the biofilm structure, we allowed the contact with the biofilm for 2 hours. Then, we washed the capillary with LB by increasing the flow rate to thoroughly rinse off any surface adhered PmB and remove surface debris due to the PmB. Fresh LB medium was then flowed for 24 hours at 37°C.

We observed that the control held up better to the capillary than the PmB-treated biofilm, suggesting that part of the biofilm treated with PmB was detached possibly due to a degradation of the surface layer. It is also noteworthy that 24 hours after the PmB1600 treatment, the biofilm successfully re-established and resumed growth (Figure 5C). However, it is different from the control as its structure is much more localized and densified, exhibiting large aggregate-like structures. This result suggests that the bacteria within the biofilm were protected from the PmB treatment.

### The increased resistance of the biofilm to high concentrations of PmB is not linked to an overexpression of matrix biofilm genes

To determine if the phenotypes observed through microscopy and crystal violet assays were a consequence of differential gene expression encoding biofilm essential components in *V. cholerae*, we performed transcriptomic analysis for genes encoding matrix proteins and VPS (Figure 6). The *vpsU* gene reports on locus I of the *vps* genes, while *vpsL* reports on locus II transcription. A1552R biofilms were allowed to grow overnight, and the medium was replaced with fresh LB containing 1600 µg/ml of PmB. The biofilms were incubated for 15 minutes or 6 hours. RNAs were extracted and retro-transcribed, and gene expression was determined by qPCR (Figure 6).

**Figure 6:**
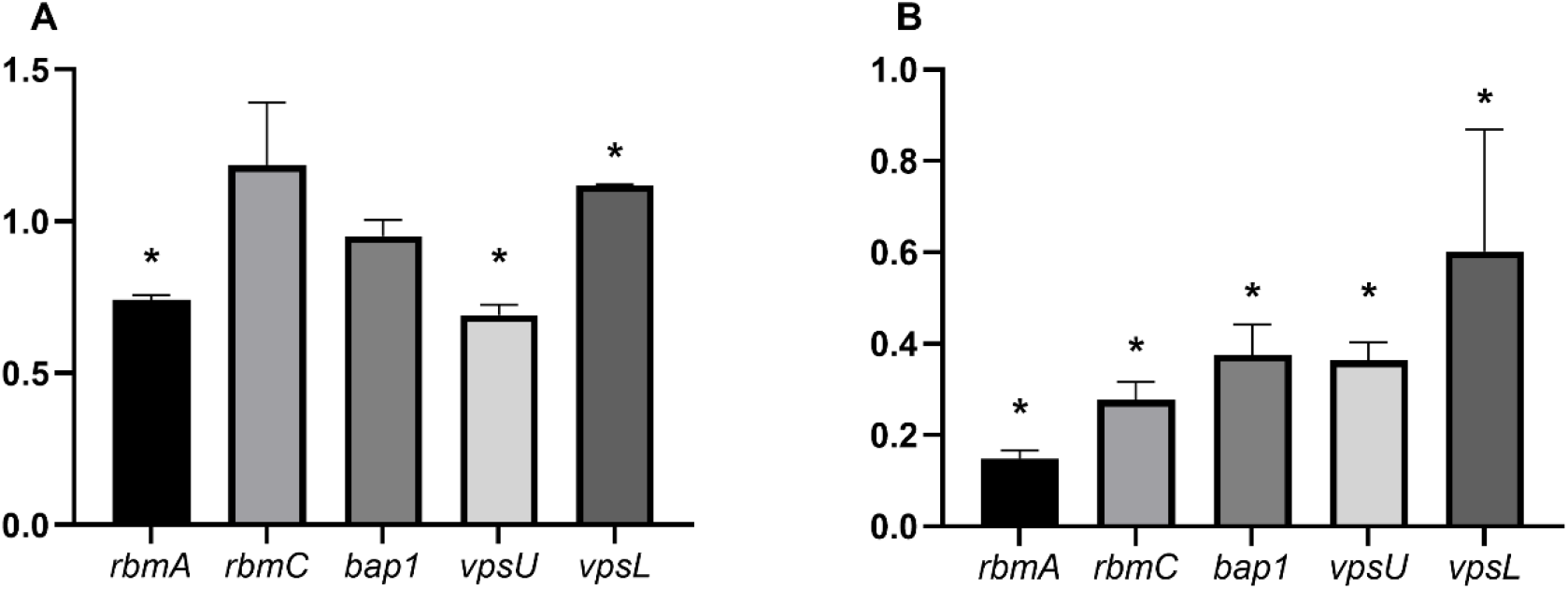
*V. cholerae*’s biofilm matrix genes’ expression after exposure to 1600 µg/ml of polymyxin B (PmB). Preformed biofilms were treated with 1600 µg/ml of PmB for 15 min (A) or 6h (B). Total RNA was extracted and retrotranscribed. The expression of *rbmA*, *rbmC*, *bap1*, *vpsU* and *vpsL* was measured by quantitative PCR in the presence of PmB in comparison to the non-treated cells, and normalized using *recA*. Data represent mean ± SD of four independent experiments conducted in triplicates. Asterisk represents a significant difference in expression between treated and non-treated cells (P<0.05). ns, non-significative difference.

Since the results from microfluidic experiments demonstrated that adaptation of biofilms to 1600 µg/ml of PmB occurred within the first 20 minutes of treatment, we attempted to observe significant differences in gene expression at 15 minutes of PmB1600 treatment. Additionally, to determine the long-term impact of PmB, we also analyzed the expression of biofilm matrix genes after 6 hours of treatment.

After 15 minutes, we observed a downregulation of *rbmA* and *vpsU* genes and a slight overexpression of *vpsL* in the PmB treated biofilm in comparison to the non-treated biofilm (Figure 6). There were no changes observed for *bap1* and *rbmC* expression in both conditions (Figure 6). However, all the genes were strongly repressed after 6 hours of PmB exposure (Figure 6). The *rbm* genes were the most severely downregulated (Figure 6). We did not observe any increase in the expression of these genes that could explain the increase of the biomass in both static and flow systems, suggesting that it is not due to an over-expression of the biofilm matrix genes.

### Treatment of the biofilm with high concentration of PmB induces a massive release of DNA

To understand the opacification mechanism of the biofilm observed in the microfluidic system, we stained the biofilms of interest with two types of DNA dyes, NucBlue™ and Propidium Iodide (PI) (Figure 7). The NucBlue stains both intracellular and extracellular nucleic acid while the PI stains only the extracellular or the nucleic acids inside severely damaged cells. The 12h preformed biofilms were treated simultaneously with PmB and fluorescent dyes. After 30 minutes of incubation, we examined the biofilms under a fluorescence microscope. The exposure was set to the channels with PmB1600, and the parameters were kept observing the control conditions. The fluorescence is more intense in the capillary with PmB compared to the non-treated controls (Figure 7). Additionally, the fluorescence appears concentrated within the biofilm aggregates in the PmB treated biofilms. These results suggest that PmB induces a massive release of DNA in response to PmB treatment, likely due to the lysis of bacteria on the biofilm surface.

**Figure 7:**
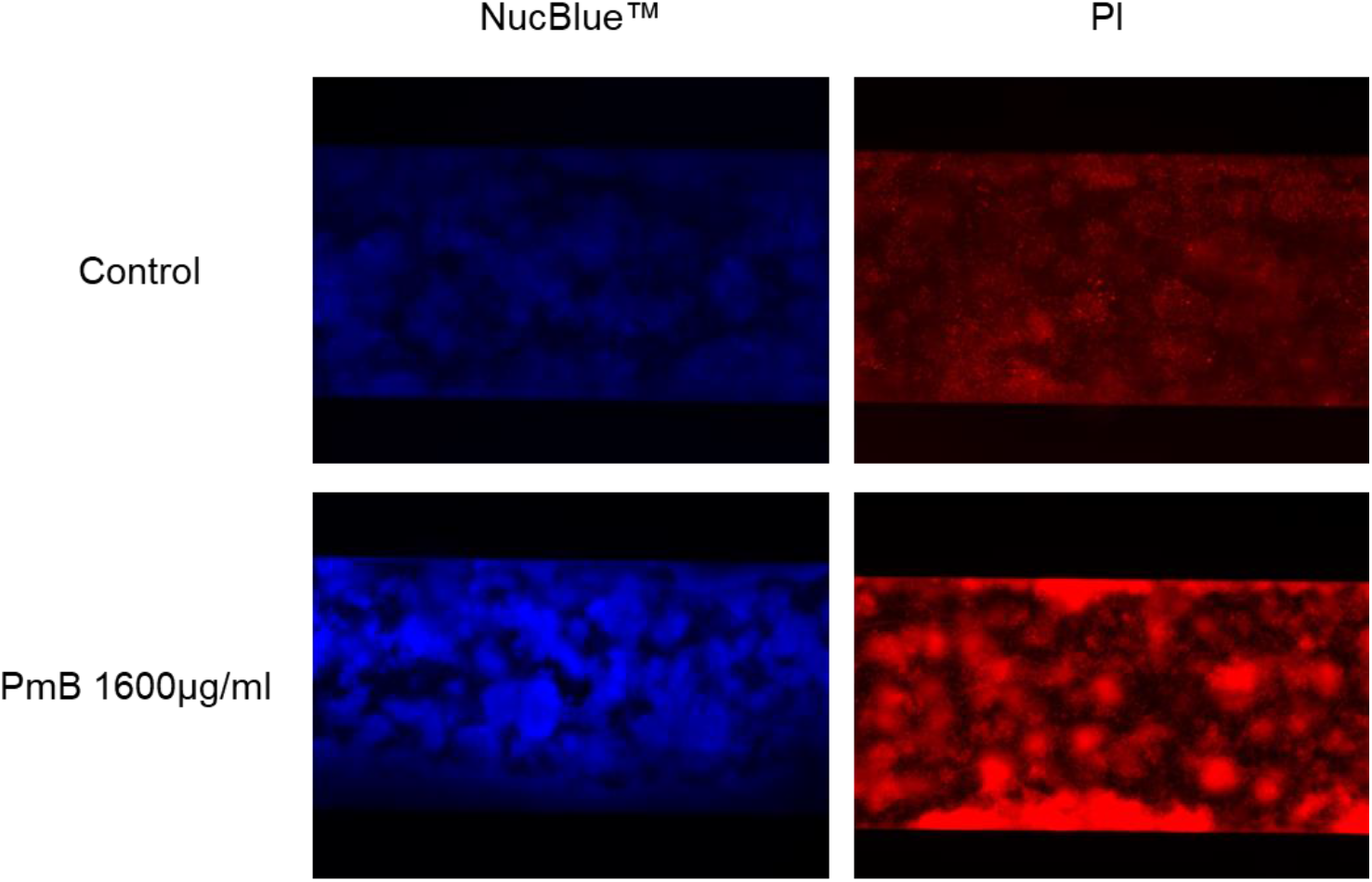
Fluorescence staining of polymyxin B (PmB)-treated biofilms. A preformed biofilm was treated for 24h with 1600 µg/ml of PmB in the microfluidic system. A staining with Nucblue^TM^ (left lower panel) or with PI (right lower panel) was performed and the fluorescence of the biofilm observed. The microscope parameters were maintained between stained and unstained conditions. Data are representative of at least 3 independent experiments.

## Discussion

In this study, we first aimed to characterize the antimicrobial resistance of *V. cholerae* under anaerobic respiration in comparison with aerobic conditions. *V. cholerae* primarily utilizes anaerobic respiration to support its metabolism during intestinal colonization, with fermentation playing a secondary role in a highly complex regulatory system (32). Therefore, a purely fermentative metabolism does not represent the reality of the infectious environment and leads to significant differences in resistance (33). To address this issue, we used a LB medium enriched with TMAO for anaerobic growth. TMAO is added at a concentration of 50 mM to replace oxygen as the final electron acceptor, ensuring anaerobic respiration (28).

Our results show that *V. cholerae* A1552 and A1552R are more resistant to streptomycin and kanamycin in anaerobiosis than in aerobiosis. These two antibiotics, from the aminoglycoside family, require an intact electron transport chain to be imported in the cell, and thus are less effective against anaerobic bacteria (34). The anaerobic-adapted electron transport chain in *V. cholerae* comprises several analogous proteins that differ from the classical chain in aerobic environments (32). Besides, the anaerobic growth with and without antimicrobials is significantly lower than aerobic growth, with an OD_600nm_ of approximately 0.1 and 0.5, respectively. This difference is not surprising, considering that oxidative respiration is the most efficient method of ATP generation (35). Furthermore, the viability of *V. cholerae* is reduced after 4 hours of growth using TMAO, due to the accumulation of toxic metabolic wastes in the medium, leading the bacteria to prematurely enter the stationary phase compared to aerobic growth (36). Thus, the impaired growth under anaerobic conditions resulting from a reduced metabolic activity might also explain the increased resistance to aminoglycosides.

We observed a difference in resistance to PmB, for which a 3-fold increase in the CI_50_ is observed under anaerobic conditions compared to aerobiosis. The main mechanisms of PmB resistance in *V. cholerae* involve alterations in LPS and vesicle production (29, 37, 38). A recent study demonstrated LPS modification under anaerobic conditions in *V. cholerae* when grown in a TMAO- and bile-enriched medium (39). Although in a different medium, it is possible that LPS modifications may occur under anaerobic conditions and could influence the sensitivity of *V. cholerae* to PmB. Given these results, we focused on PmB to further characterize the impact of anaerobic respiration on *V. cholerae* resistance.

An important part of antimicrobial resistance is due to biofilm formation and the biofilms of *V. cholerae* are important for environmental survival and pathogenicity (14, 40, 41). Thus, we investigated the impact of the oxygen availability and PmB on biofilm formation in *V. cholerae.* A lower biofilm production in the anaerobic environment was observed, which is consistent with the reduced growth under anaerobic in comparison to aerobic conditions. A previous study also drawn similar conclusions, stating that the hypoxic environment limits biofilm formation in *V. cholerae* compared to an oxygen-present environment (42). Besides, we observed that biofilm production is inhibited from 25 µg/ml of PmB in all strains under aerobic conditions. These findings align with our previous study, attributing the decrease in biofilm formation to the loss of the flagellum, an important adhesion element in *V. cholerae* (30, 43). Thus, subinhibitory concentrations of PmB induce a decrease in biofilm adhesion to its substrate, resulting in the decrease in biomass production. In anaerobiosis, biofilm production is inhibited from 3 µg/ml, which is 75 times lower than the anaerobic MIC, while our results show that the planktonic bacteria are more resistant under anaerobic growth. It has been recently demonstrated that the two-component system ArcAB, a key system for the adaptation to oxygen, is negatively regulating the flagellum’s genes expression and motility (44). In low oxygen conditions, *arcA* is strongly upregulated, leading to a significant reduction of the flagellum production (44). Thus, the negative effect of PmB on the flagellum, and subsequently on biofilm formation, might be enhanced by the increased expression of *arcA* under anaerobic growth.

After determining the impact of PmB on biofilm formation, we investigated its effect on mature 24-hour biofilms. We first observed that there is a stimulation of the biofilm formation at low concentrations (bellow 25 µg/ml of PmB) for the rough strain A1552R in the presence of oxygen.

This phenotype is not observed in anaerobiosis. It has been shown in other bacteria that sub-inhibitory concentrations of antibiotics stimulate the biofilm production (45). Then, we observed a dose-dependent effect of PmB on the biofilm with almost 50% of the biomass reduction both in the presence and in absence of oxygen. This effect occurs at lower concentrations in anaerobiosis than in aerobiosis, which might be explained by the lower initial biomass of the biofilm.

Intriguingly, at high concentrations of PmB (> 400 µg/ml), we observed a higher biofilm biomass than at lower concentrations, especially under aerobic conditions. We also observed a 3- dimensional mushroom-like structure of the bacterial colonies on solid media, suggesting that the bacteria are growing on top of each other to protect themselves from the PmB contained in the media. With a microfluidic device, we assessed the biofilm response to PmB over time. A rapid opacification of the biofilm was observed within 20 minutes of PmB1600 exposure. An analysis of the essential matrix genes’ expression after 15 min of PmB1600 treatment showed that it is not due to an over-expression of these components. Our next hypothesis was that the increased opacity of the biofilm could be due to the accumulation of DNA generated by the lysis of many surface bacteria. To investigate this possibility, we stained the biofilms with PI and Nucblue^TM^, two fluorescent DNA labelling dyes, and showed that the biofilms treated with PmB1600 fluoresce much more than the untreated control. These results suggest that a large amount of DNA is present on the surface of the biofilms treated with PmB1600. Since the biofilm remains intact 24 hours after treatment with PmB1600, unlike after the treatment with PmB200, it is likely that this protection requires a high concentration of PmB to release sufficient DNA to envelope and protect the biofilm. The protective characteristics of DNA are not a new concept. DNA is known to be an adhesive molecule that participates in the biofilm structure by linking polymers, proteins, and other components of the matrix to stabilize the structure (46). The polarity of DNA is also described to sequester some antibiotics like aminoglycosides and cationic antimicrobials, such as PmB (47). Therefore, we propose a model in which DNA is released following bacterial lysis by high concentrations of PmB. This DNA may sequester the PmB and even reinforce the biofilm structure, thereby protecting the bacteria inside the biofilm by acting as a shield.

The extreme resistance of biofilms to antibiotics is well-known; however, several authors consider AMPs as potent anti-biofilm agents due to their action on bacterial membranes (48, 49, 50). AMPs are interesting because they can act on persisting bacteria in addition to metabolically active bacteria, unlike conventional antibiotics (50, 51). The action of AMPs as anti-biofilm agents is highly effective in the initial stages of biofilm formation, as demonstrated by our previous work and others (30, 48, 52). However, for the AMPs to have a bactericidal effect, they must reach the target bacteria, which may not be possible depending on the complexity and maturity of the biofilm. The action of AMPs on the extracellular matrix is poorly understood, and some components of the matrix, such as nucleic acids, may have inhibitory effects on AMPs, limiting their action on already-formed biofilms (52).

## Author contribution

Conceptualization: J.P.F. and M.D.; Data curation: J.P.F. and M.D.; Formal analysis, J.P.F. and M.D.; Investigation: J.P.F.; Methodology: J.P.F.; Validation: J.P.F. and M.D.; Visualization: J.P.F. Writing – original draft: J.P.F. and M.D. Writing – review and editing: all authors. Resources: M.D.; Supervision: M.D.; Funding acquisition: M.D.; Project administration, M.D.

## Funding

This work was supported by the Natural Sciences and Engineering Research Council of Canada (NSERC; http://www.nserc-crsng.gc.ca/index_eng.asp) Discovery grant number RGPIN-2017- 05322 (Principal Investigator: MD). J.P.F. received financial support from the FRQNT scholarship program, from the NSERC’s Canada Graduate Scholarships Master’s program (CGS M) and the J.A. DeSève Scholarship from the Graduate and Postdoctoral Studies of the University of Montreal (ESP).

## Competing interests statement

The authors declare there are no competing interests.

## Data availability statement

Data generated or analyzed during this study are provided in full within the published article and its supplementary materials.

